# GIMMEcpg: Global Imputation of Mean CpG MEthylation in Real-time

**DOI:** 10.1101/2025.03.17.643746

**Authors:** Ismail Moghul, Niuzheng Chai, Nikolas Pontikos, Alison Hardcastle, Javier Herrero, Stephan Beck

## Abstract

Whole-genome DNA methylation (methylome) analysis is of broad interest to biomedical research due to its central role in human development and disease. However, generating high-quality methylomes at scale remains challenging due to inherent technical limitations. While imputation has the potential to help overcome this problem, no existing approach adequately addresses the scaling issue.

Here, we present GIMMEcpg (Global Imputation of Mean cpg MEthylation), a novel imputation tool that scales efficiently from single samples to large cohort studies. GIMMEcpg uses a custom feature dataset built from known CpG sites within the same dataset to impute missing values by calculating the distance-weighted mean of the methylation value of the two immediately neighbouring CpG sites.

We benchmarked GIMMEcpg for speed and accuracy against multiple imputation methods using downsampled datasets produced from high-quality (∼100x) Whole Genome Bisulfite Sequencing (WGBS) data. With a 10x downsampled dataset, GIMMEcpg was able to process the dataset and impute 9.14 Million CpG sites within 7 seconds (R: 0.78, MAE: +5.6%, RMSE: +10.9%). Our results demonstrate that GIMMEcpg is 39-2,562 times faster than three existing methylation imputation tools (BoostMe, DeepCpG, and MethImpute) while maintaining comparable accuracy.

To quantify GIMMEcpg’s scalability, we applied it to the most extensive single collection of WGBS data (N=645 at variable coverage) from the EpiATLAS generated by the International Human Epigenome Consortium (IHEC). Using a single, standard CPU server, GIMMEcpg processed and imputed an additional 2.4 billion CpG methylation values across the 645 datasets in less than a day, enriching the EpiATLAS methylome resource by 20%. This demonstrates that GIMMEcpg scales to large cohort studies with only a subtle impact on accuracy, as illustrated by our benchmark.

We also developed a machine learning variant, GIMMEcpg.ml, which delivers a higher accuracy compared to existing methodologies. Using the same 10x downsampled benchmarking dataset, GIMMEcpg.ml achieved a Person Correlation of 0.87 compared to the ground truth, representing an improvement of 0.11 over the best performing alternative method. Additionally, GIMMEcpg.ml has a Mean Absolute Error (MAE) of 8.67%, which is 2.63% lower than the most accurate performing alternative. While this enhanced accuracy comes at the cost of increased computation requirements, GIMMEcpg.ml is a useful tool where higher accuracy is preferred over scalability.

For increased accessibility, GIMMEcpg is freely available under an MIT license as R and Python packages at https://github.com/ucl-medical-genomics/gimmecpg-r and https://github.com/ucl-medical-genomics/gimmecpg-python, respectively.

## Introduction

Over the past decades, plummeting sequencing costs have made genomic and epigenomic analysis more accessible to researchers (Kris A. Wetterstran n.d.; Heidi Chial 2008). As a result, whole-genome DNA methylation (methylome) analysis has become possible (Beck 2010) and has been applied to many studies in health and disease (Day & Sweatt 2010; Jin & Liu 2018; Feinberg 2007). However, the vast majority of existing methylome data is still based on array technologies that only cover 1-3% of the human methylome. In fact, over 99% of studies on the EWAS atlas were carried out using arrays (Li et al. 2019). Sequencing-based methylome analysis first gained popularity with the introduction of Reduced Representation Bisulfite Sequencing (RRBS) (Meissner et al. 2005) and targeted bisulfite sequencing technologies, but these typically still only cover 14-15% of the entire methylome (Tanić et al. 2022). In comparison, only a few studies to date involved large-scale Whole Genome Bisulfite Sequencing (WGBS), most notably the DNA Methylation Atlas (Loyfer et al. 2023) and the IHEC EpiATLAS (EpiATLAS Consortium 2025).

Despite its limitations, bisulfite treatment has been the gold standard for the analysis of 5-methylcytosine (5mC) for over 30 years (Frommer et al. 1992). However, it is important to note that this method cannot distinguish 5mC from its oxidised forms 5-hydroxymethylcytosine (5hmC), 5-formylcytosine (5fC) and 5-carboxycytosine (5caC) (Wu & Zhang 2017). Therefore, it can only distinguish between modified and unmodified cytosines. For the purpose of this study, we designate all modified cytosines as ‘methylated’, as the vast majority of modifications constitute 5mC. Although bisulfite sequencing remains popular, many alternative and potentially superior methods have been developed for mapping DNA methylation and other epigenetic modifications by sequencing technologies (Chen et al. 2023).

Bisulfite treatment also introduces multiple technical challenges that complicate the analysis. One significant issue is incomplete conversion, where some unmethylated cytosines are not converted into uracil, making it impossible to differentiate between a methylated cytosine and an unconverted, unmethylated cytosine. The bisulfite conversion rate typically ranges from 95-98% (Warnecke et al. 2002). In human WGBS datasets, this means that approximately 1.4 million cytosines are unconverted (assuming a total of 29 million haploid CpGs and a 95% conversion rate), which will, as a result, be incorrectly interpreted as methylated. Another challenge is DNA fragmentation, as bisulfite treatment causes extensive DNA fragmentation (Simpson et al. 2017), negatively affecting downstream PCR amplification.

Furthermore, these issues are particularly relevant in the context of bulk-cell sequencing. Bulk cell samples typically contain multiple cell types of varied developmental stages, which affects the precision and accuracy of calculating average CpG methylation values. It is, therefore, crucial to ensure sufficient sequence coverage over each CpG site to accurately call methylation values and, where possible, deconvolve them into their constituent cell types using a cell type deconvolution algorithm (reviewed in (Teschendorff & Relton 2018).

Imputation can overcome some of these limitations and has been used in genetic studies for several decades with high accuracy. It is now a standard part of many Genome-Wide Association Studies (GWAS) and other genomic analyses (Marchini & Howie 2010). Over the last few years, imputation has also been explored in the context of DNA methylation. Several tools have been developed for this purpose, each with its own approach and scope.

DeepCpG, initially designed for single-cell WGBS data, has also shown applicability to bulk-cell WGBS data. It utilises deep learning techniques to determine associations between DNA sequence patterns (1kb) and between neighbouring CpG sites (50 CpG sites) (Angermueller et al. 2017). Melissa, another tool developed for single-cell data, utilises a Bayesian hierarchical method to cluster local methylation patterns and leverages this information to impute missing data effectively (Kapourani & Sanguinetti 2019).

For bulk-cell WGBS data, tools like BoostMe and METHimpute have been developed. BoostMe uses a gradient boosting algorithm to extract useful information from the two immediate, neighbouring CpG sites as well as the sample average (Zou et al. 2018). On the other hand, METHimpute leverages a Hidden Markov Model-based binomial test that learns from the methylation status of neighbouring cytosines (Taudt et al. 2018).

ChromImpute is the first imputation method that was designed for large epigenomic studies with multi-dimensional data types, including histone modifications and RNA-seq (Ernst & Kellis 2015). It can predict the methylation signal in a target sample based on two classes of features: signal from other data types in the same sample and the corresponding methylation signal in other samples. The ability of ChromImpute to integrate information across multiple dataset types (including histone modifications, RNA-Seq, and WGBS) and across multiple samples made it the method of choice for imputation of major epigenomic resources such as EpiMap (Boix et al. 2021) and the IHEC EpiATLAS (EpiATLAS Consortium 2025). Even though ChromImpute can impute WGBS if other histone and/or RNA-seq data are present, it is unable to do so with just WGBS data which is the focus of this study. For this reason, ChromImpute was excluded from the benchmarking presented here.

While these imputation tools are valuable for extracting maximum value from a dataset, many come with high computational demands that limit their scalability to large datasets. Addressing this gap, we introduce GIMMEcpg (Global Imputation of Mean CpG Methylation), a novel imputation tool specifically designed to efficiently scale from analysing a single sample to handling extensive cohort studies.

## Results

### GIMMEcpg: Global Imputation of Mean CpG Methylation

GIMMEcpg is built upon the established principle that the methylation value of CpG sites correlates highly with neighbouring CpG sites, with this correlation increasing as the distance between the neighbouring CpG sites decreases (Eckhardt et al. 2006; Moghul 2021; Li et al. 2010). This key feature has also been leveraged by other imputation tools, such as DeepCpG and BoostMe, for imputing missing values.

Eckhardt et al. (2006) first demonstrated that a significant correlation was observed between neighbouring CpG sites up to distances of 1Kb, which then rapidly deteriorated for distances larger than 2kb (Eckhardt et al. 2006). However, as this study was carried out using old Amplicon-based technology over 15 years ago, we decided to re-evaluate this relationship using high-quality (100x coverage) WGBS datasets.

Our analysis, as illustrated in Supplementary Figure 1, reveals that the Pearson correlation between neighbouring CpG sites decreases more rapidly with increasing genomic distance than previously reported. At very short distances, the Pearson correlation approaches 1, but it rapidly declines linearly to a correlation of 0.3 at 250bp. Beyond this distance, although some correlation persists between neighbouring CpG sites, it is considerably weaker, reaching random levels of correlation past 1.5kb.

GIMMEcpg builds on this information by employing a simple yet effective imputation strategy, where the distance-weighted mean of the two neighbouring CpG sites is calculated, as described in the equation below. This simplified mathematical approach significantly reduces computational requirements compared to more complex Machine Learning or Deep Learning models while maintaining high accuracy within biologically relevant distances (i.e. below 1kb).

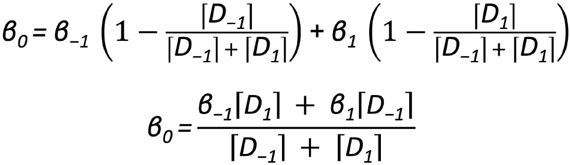

***Equation 1***: *The methylation value (β*_*0*_*) of a CpG site with an unknown methylation value can be defined as the weighted average of the methylation value of the neighbouring CpG sites. The methylation value of the upstream and downstream neighbouring CpG sites are denoted by β*_*1*_ *and β*_*-1*_, *respectively. The linear distance in base pairs between the focal CpG site (with the unknown methylation value) and each downstream and upstream neighbouring CpG site are denoted by D*_*1*_ *and D*_*-1*_, *respectively*.

This formula is based on the assumption that there is a correlation between neighbouring CpG sites. Therefore, it is important to avoid imputing CpG sites using this formula where this assumption no longer holds. By default, GIMMEcpg does not impute CpG sites if both neighbouring CpG sites are further than 1KB away from the focal CpG site.

GIMMEcpg provides the distance to the closest neighbouring CpG as a proxy for a confidence value to aid in interpretation and allow for customised downstream analysis. For example, end-users are able to choose a stricter distance cut-off if desired, balancing accuracy with the number of CpG sites imputed based on their specific research needs.

### Baseline Accuracy

Establishing a reliable ground truth dataset is crucial to assessing the performance of imputation methods like GIMMEcpg. Therefore, we utilised high-quality (100x coverage) bulk-cell WGBS data from the IHEC EpiATLAS for this purpose (EpiATLAS Consortium 2025). Our approach involved downsampling two high-coverage WGBS datasets to simulate WGBS datasets with lower CpG coverage. This process resulted in 48 downsampled datasets, comprising six datasets at eight different coverage levels. In this manner, the actual methylation value is known for each missing data point in the downsampled dataset, allowing for precise accuracy assessment of any subsequent imputation.

It is essential to note that the imputation accuracy calculated using a lower coverage downsampled dataset compared to a higher coverage original dataset is likely to be overestimated. This overestimation occurs due to an expected decrease in accuracy from having data with a lower CpG coverage. To account for this, it is important to determine the baseline error (or residual noise) by comparing methylation values in each downsampled dataset to the original dataset for CpG sites present before imputation (i.e., the baseline accuracy).

Our analysis revealed that at a 10x coverage, a very high Pearson correlation of 0.96 exists between the downsampled dataset and the corresponding original dataset. However, when examining the Mean Absolute Error (MAE) and the Root Mean Squared Error (RMSE), it becomes clear that some additional variance (or error) was introduced as a result of downsampling. These effects appear relatively small, with an MAE of 6% and RMSE of 8.8%. The baseline accuracy present in the datasets downsampled to a 10x coverage is shown in Figure 1A,D,E, while the baseline accuracy for all the downsampled datasets with coverages ranging from 5x to 60x are shown in Supplementary Figure 2.

**Figure 1:**
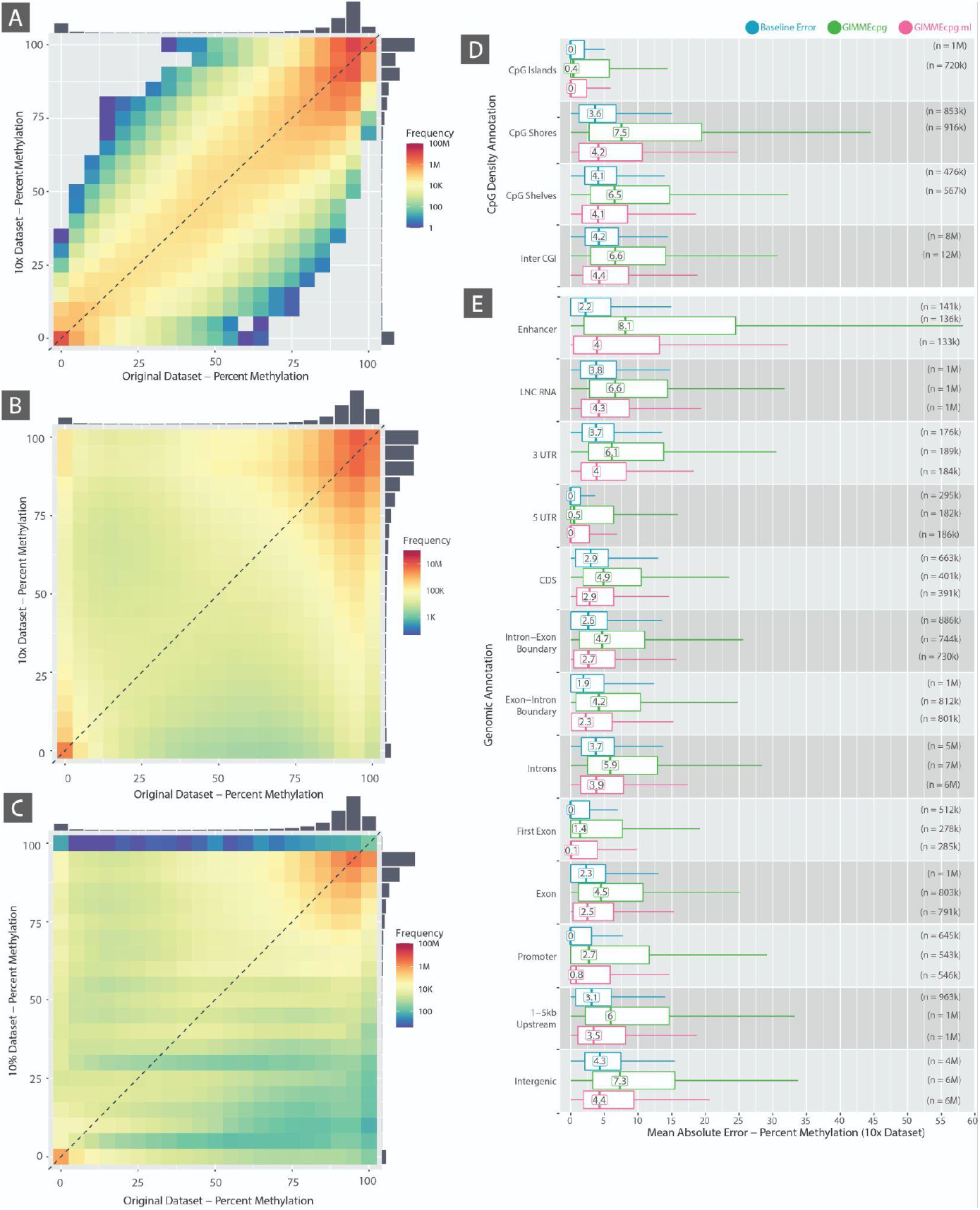
[A]: Baseline error in downsampled datasets with a 10x coverage. (i.e. residual noise in the dataset due to being at lower coverage). A distribution, averaged over six downsampled datasets with a 10x coverage, comparing the methylation value in the downsampled dataset and the original 100x methylation values used here as ground truth. The plot only includes information from CpG sites with a greater than a 10x read depth. Any deviation from the y = x-axis corresponds to an error in the methylation value. The baseline accuracy for all the downsampled datasets with coverages ranging from 5x to 60x are shown in Supplementary Figure 2. **[B] GIMMEcpg Imputation Accuracy in downsampled datasets with 10x coverage.** A distribution of imputed values, comparing the imputed methylation value and the corresponding actual methylation value in the original dataset. Any deviation from the y = x-axis corresponds to an error in the methylation value. However, it is important to note that these plots do not take into account the baseline error and, consequently, are expected to be overestimated. The GIMMEcpg Imputation accuracy for all the downsampled datasets with coverages ranging from 5x to 60x are shown in Supplementary Figure 3. **[C] GIMMEcpg.ml Imputation Accuracy in downsampled datasets with 10x coverage**. A distribution of imputed values, comparing the imputed methylation value and the corresponding actual methylation value in the original dataset. Any deviation from the y = x-axis corresponds to an error in the methylation value. However, it is important to note that these plots do not take into account the baseline error and, consequently, are expected to be overestimated. The GIMMEcpg.ml Imputation accuracy for all the downsampled datasets with coverages ranging from 5x to 60x are shown in Supplementary Figure 4. **[D-E]** Box plots showing the distribution of the baseline absolute error (green and the error associated with GIMMEcpg (blue) and GIMMEcpg.ml (pink), grouped according to the CpG density and the genomic annotation of the CpG site. Outliers are not shown on the plot due to the number of outliers; rather, the box-plot whiskers are extended to the maxima and minima with a dashed line. The box plots show summarised information from six downsampled datasets with a 10x coverage.

Interestingly, when grouping the CpG sites based on genomic features, the MAE is not equivalent across different genomic features despite the coverage downsampling being carried out randomly. Furthermore, the MAE is minimal in certain genomic features such as CpG islands, 5’ UTRs, promoters and first exons, while being elevated in other genomic features like intergenic regions, introns, 3’ UTRs and LNC RNA.

These results also highlight that even at 30x, there is an MAE of 3.3% and an RMSE of 5%. This finding highlights that each methylation value is only accurate to ±3%, compared to when sequenced at 100x. The implication of this is significant in that 30x WGBS should not be used to infer DMPs with a differential methylation value smaller than 6%.

### GIMMEcpg Performance

GIMMEcpg was able to impute missing data in a WGBS dataset with 10x coverage in 7 seconds while using under 40GB of RAM. In these six downsampled datasets with a 10x coverage, an additional 9.1 million CpG sites were imputed in each dataset, with an average Pearson correlation of 0.78, an MAE of 11.6% and an RMSE of 19.7%. These numbers are slightly higher than the baseline error in the same dataset (i.e., an increased MAE of +5.6% and an increased RMSE of +10.9%).

These results suggest that GIMMEcpg is able to process and impute a complete WGBS dataset extremely efficiently while ensuring a relatively high level of accuracy. The performance of GIMMEcpg when imputing the downsampled dataset with a 10x coverage is shown in Figure 1B,D,E, while the performance of GIMMEcpg for all the downsampled datasets with coverages ranging from 5x to 60x are shown in Supplementary Figure 3.

Furthermore, the accuracy of the imputed values can be further explored by separating the imputed CpG sites by the genomic annotation. The imputed methylation value has a fairly low MAE in CpG Islands (0.4% at 10x), 5’ UTRs (0.5% at 10x), and first exons (1.4%). These results can be explained by the fact that these regions are more CpG-dense regions, where GIMMEcpg works best (i.e., regions where the neighbouring CpG site is close by).

### GIMMEcpg.ml

An alternative machine learning approach using model stacking was also explored, where existing data from within the same dataset were utilised to train multiple models which are then merged to produce a single ensemble model. As part of this approach, multiple machine learning techniques, including XgBoost (Chen & Guestrin 2016), Gradient Boosting Machines (GBM) (Friedman 2001), Extremely Randomised Trees (XRT) (Geurts et al. 2006), Distributed Random Forests (DRF) (Cevid et al. 2022), Generalised Linear Models (GLM) (Nelder & Wedderburn 1972) and a fully-connected, multi-layer Artificial Neural Networks (ANN) (Hopfield 1988) were employed. Furthermore, an automated grid search was utilised to improve the model’s accuracy through 5-fold cross-validation, before being merged into a single ensemble model.

The feature dataset is generated from known data (i.e. CpG sites with a corresponding methylation value) within the dataset. The feature dataset is inspired from information utilised within the GIMMEcpg approach. Here, the methylation value of known CpG sites is used as a feature, together with the methylation value and distance to the two immediately neighbouring CpG sites.

In this manner, a dataset with five features is produced (Figure 2B) containing the focal methylation value, together with the genomic distance and the methylation value of the two immediately neighbouring CpG sites. After training, the best-performing model from cross-validation is selected and used to impute the DNA methylation value of missing CpG sites using a similar dataset generated for the missing CpG sites.

**Figure 2:**
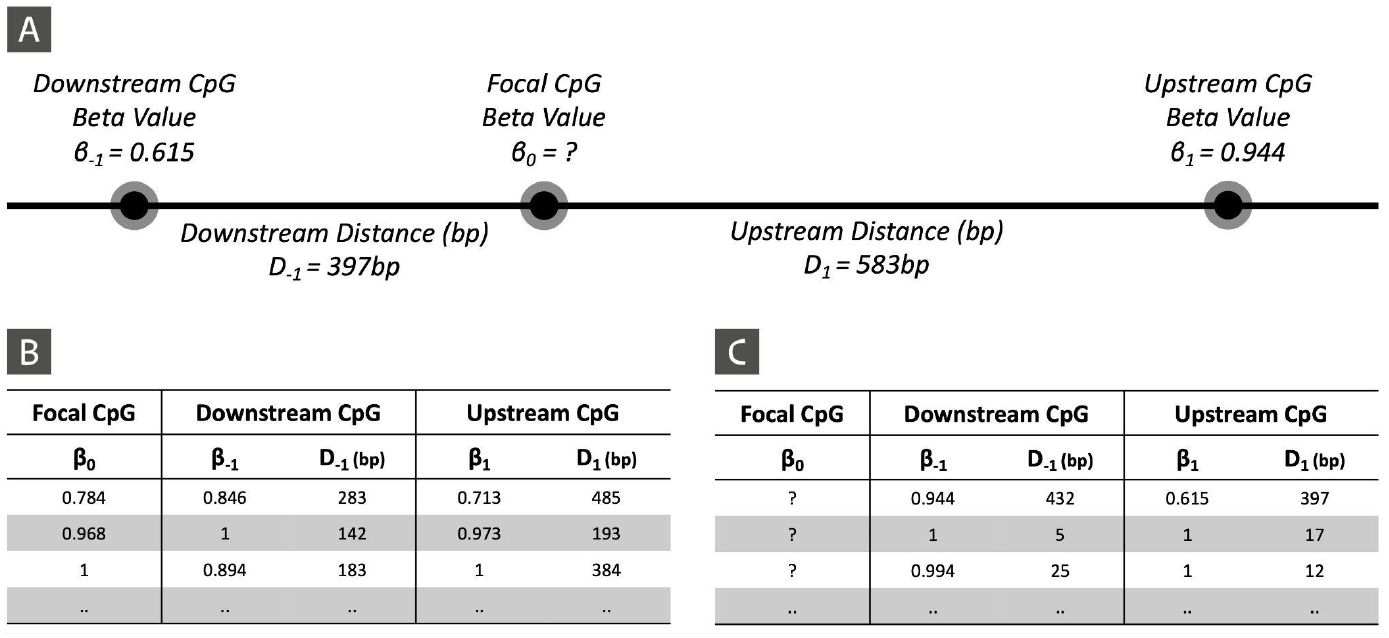
Imputation using feature dataset produced from information on immediate CpG neighbours. **[A]** A schematic showing a CpG site (β_0_) with an unknown methylation value and its two immediate neighbouring CpG sites. The methylation value of the upstream and downstream neighbouring CpG sites are denoted by β_1_ and β_-1_, respectively. The linear distance in base pairs between the focal CpG site (with the unknown methylation value) and each downstream and upstream neighbouring CpG site are denoted by D_1_ and D_-1_, respectively. The linear distance in base pairs between the focal CpG site (with the unknown methylation value) and each downstream and upstream neighbouring CpG site are denoted by C_1_ and C_-1_, respectively. **[B]** The feature table shows cases where the focal CpG site has a known methylation value. This would be used to train the model, where the algorithm will attempt to calculate a value close to the actual methylation value using the rest of the data in the feature dataset. **[C]** For each CpG site with an unknown methylation value, a similar feature dataset is generated with the β_0_ value left blank. A trained model can then be used to calculate the value of β_0_ using the information in the remainder of the feature dataset.

When applied to the six downsampled datasets at a coverage of 10x and given 30 minutes of training time, GIMMEcpg.ml imputed an additional 13.8 million CpG sites at an MAE of 8.67% and a Pearson correlation of 0.87 (See Figure 1C,D,E and Supplementary Figure 4). These results demonstrate that GIMMEcpg.ml is able to successfully train a highly accurate model from data within the same sample. Furthermore, by exploring the methylation value through grouping by genomic annotation, it becomes apparent that GIMMEcpg.ml is significantly more accurate than GIMMEcpg and works best in CpG-dense regions such as CpG Islands, 5 UTRs, first exons, and promoters.

These results suggest that the machine learning approach of GIMMEcpg.ml offers a significant improvement in accuracy over the standard GIMMEcpg method. However, this comes at the cost of increased computational time and complexity. The choice between GIMMEcpg and GIMMEcpg.ml may depend on the specific requirements of a research project, balancing the trade-off between imputation accuracy, computational efficiency and scalability.

### Benchmarking against alternative methods

We systematically evaluated and benchmarked purpose-built DNA methylation imputation algorithms (such as DeepCpG, BoostME and MethImpute) together with mean imputation as a baseline imputation methodology. A total of ten different methodologies, also summarised in Table 1, were applied to the 48 downsampled datasets (described above), and the imputation accuracy (MAE, RMSE, R) and compute performance (time taken and memory utilised) were calculated. The accuracy and performance results are shown in Table 1 and Figure 3.

**Table 1.** Accuracy metrics for GIMMEcpg and existing imputation methodologies at eight different coverage levels ranging from 5x to 60x. R: Pearson Correlation; MAE: Mean Absolute Error (%), RMSE: Root Mean Squared Error (%), RAM - Memory used (GB), Time (seconds). The colour scale is relative to each coverage block and is described in the key, where green is shown for better-performing values (i.e. higher number of CpGs or higher R, or lower error rates), and red used for worse values.

| Imputation Method | D05 |  |  |  |  |  | D07 |  |  |  |  |  | D10 |  |  |  |  |  |
| --- | --- | --- | --- | --- | --- | --- | --- | --- | --- | --- | --- | --- | --- | --- | --- | --- | --- | --- |
|  | # of CpGs | R | MAE | RMSE | Time (s) | Memory | # of CpGs | R | MAE | RMSE | Time (s) | Memory | # of CpGs | R | MAE | RMSE | Time (s) | Memory |
| Baseline | 13.66 M | 0.94 | 7.9 | 11.5 | - | - | 15.97 M | 0.95 | 7 | 10.2 | - | - | 18.03 M | 0.96 | 6 | 8.8 | - | - |
| Mean | 26.03 M | - | 24.1 | 30.3 | 2 | 5.77 | 22.58 M | - | 23.2 | 29.5 | 2 | 6 | 17.68 M | - | 21.9 | 28.2 | 2 | 7 |
| MethImpute | 26.03 M | 0.7 | 18.2 | 24.7 | 708 | 601.82 | 22.58 M | 0.72 | 16.8 | 23.2 | 612 | 601 | 17.68 M | 0.71 | 16.4 | 22.5 | 684 | 601 |
| DeepCpG | 26.03 M | - | 22.9 | 30.2 | 16980 | - | 22.58 M | 0.69 | 12.4 | 22.4 | 17580 | - | 17.68 M | 0.7 | 11.6 | 21.2 | 17940 | - |
| BoostMe | - | - | - | - | - | - | 21.31 M | 0.75 | 12 | 19.5 | 276 | 182 | 16.6 M | 0.76 | 11.3 | 18.3 | 271 | 179 |
| GIMMEcpg | 4.95 M | 0.84 | 12.2 | 20.6 | 6 | 27.75 | 9.22 M | 0.82 | 11.8 | 20.1 | 7 | 33 | 9.14 M | 0.78 | 11.6 | 19.7 | 7 | 34 |
| GIMMEcpg.ml | 19.7 M | 0.88 | 8.86 | 15.2 | 1800 | 27.83 | 17.6 M | 0.88 | 8.82 | 14.9 | 1800 | 38 | 13.8 M | 0.87 | 8.67 | 14.5 | 1800 | 53 |

| Imputation Method | D15 |  |  |  |  |  | D20 |  |  |  |  |  | D25 |  |  |  |  |  |
| --- | --- | --- | --- | --- | --- | --- | --- | --- | --- | --- | --- | --- | --- | --- | --- | --- | --- | --- |
|  | # of CpGs | R | MAE | RMSE | Time (s) | Memory | # of CpGs | R | MAE | RMSE | Time (s) | Memory | # of CpGs | R | MAE | RMSE | Time (s) | Memory |
| Baseline | 19.89 M | 0.97 | 5 | 7.3 | - | - | 20.94 M | 0.98 | 4.2 | 6.3 | - | - | 21.63 M | 0.98 | 3.7 | 5.6 | - | - |
| Mean | 12.68 M | - | 20.5 | 26.9 | 2 | 8.44 | 10.09 M | - | 19.9 | 26.5 | 3 | 9 | 8.57 M | - | 19.6 | 26.3 | 3 | 9 |
| MethImpute | 12.68 M | 0.68 | 16.6 | 22.5 | 648 | 601.03 | 10.09 M | 0.66 | 17 | 22.6 | 606 | 601 | 8.57 M | 0.65 | 17.2 | 22.7 | 648 | 601 |
| DeepCpG | 12.68 M | 0.69 | 11.3 | 20.5 | 17580 | - | 10.09 M | 0.68 | 11.5 | 20.6 | 17760 | - | 8.57 M | 0.67 | 11.5 | 20.5 | 17760 | - |
| BoostMe | 11.81 M | 0.74 | 11.2 | 18 | 286 | 175.24 | 9.33 M | 0.73 | 11.3 | 18.1 | 264 | 174 | 7.88 M | 0.72 | 11.4 | 18.3 | 275 | 173 |
| GIMMEcpg | 6.26 M | 0.73 | 11.6 | 19.7 | 7 | 45.62 | 4.4 M | 0.7 | 11.8 | 20 | 7 | 46 | 3.31 M | 0.68 | 12.1 | 20.3 | 7 | 50 |
| GIMMEcpg.ml | 9.69 M | 0.84 | 9.24 | 15 | 1800 | 58.85 | 7.53 M | 0.83 | 9.45 | 15.3 | 1800 | 53 | 6.28 M | 0.82 | 9.55 | 15.4 | 1800 | 53 |

| Imputation Method | D30 |  |  |  |  |  | D60 |  |  |  |  |  |
| --- | --- | --- | --- | --- | --- | --- | --- | --- | --- | --- | --- | --- |
|  | # of CpGs | R | MAE | RMSE | Time (s) | Memory | # of CpGs | R | MAE | RMSE | Time (s) | Memory |
| Baseline | 22.12 M | 0.99 | 3.3 | 5 | - | - | 23.58 M | 1 | 1.8 | 2.8 | - | - |
| Mean | 7.59 M | - | 19.5 | 26.2 | 3 | 9.46 | 5.21 M | - | 19.5 | 26.2 | 2 | 10 |
| MethImpute | 7.59 M | 0.65 | 17.3 | 22.7 | 587 | 600.94 | 5.21 M | 0.67 | 16.7 | 21.7 | 463 | 601 |
| DeepCpG | 7.59 M | 0.67 | 11.7 | 20.6 | 17100 | - | 5.21 M | 0.64 | 12.5 | 21.2 | 17940 | - |
| BoostMe | 6.95 M | 0.71 | 11.5 | 18.4 | 278 | 172.33 | 4.69 M | 0.68 | 12.3 | 19.1 | 330 | 171 |
| GIMMEcpg | 2.65 M | 0.67 | 12.3 | 20.6 | 7 | 51.14 | 1.27 M | 0.62 | 13.1 | 21.7 | 7 | 51 |
| GIMMEcpg.ml | 5.48 M | 0.82 | 9.68 | 15.5 | 1800 | 55.43 | 3.56 M | 0.83 | 9.13 | 14.5 | 1800 | 55 |

Relative Scale for # of CpG and R
Min Max
Relative Scale for MAE, RMSE, Time, RAM

**Table 2.**
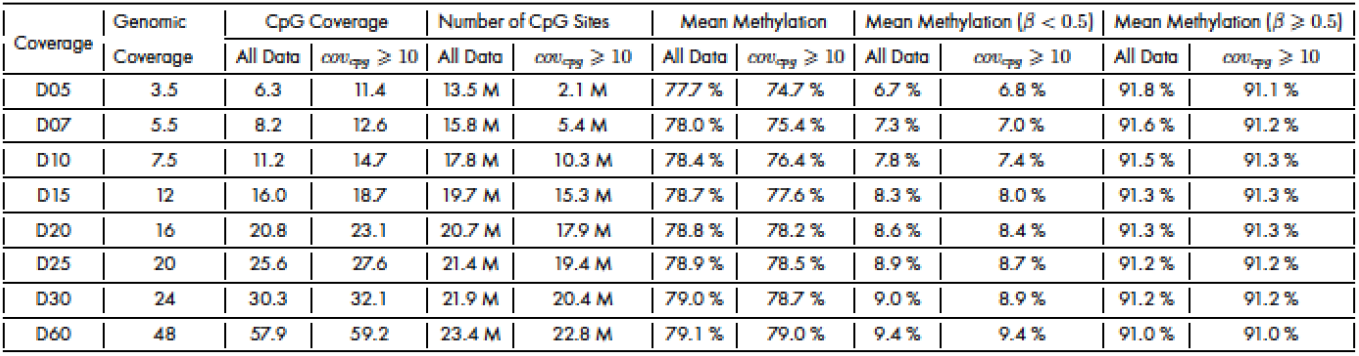
Summary of downsampled datasets produced from S1_W1 and S2_W1. The statistics have been produced by calculating a mean across six datasets at each coverage level. At the coverage level, three datasets were downsampled from S1_W1 and a further three datasets from S2_W1. Most statistics are displayed for the entire dataset (i.e. CpG sites with a coverage higher than 1) as well as for filtered subsets where the CpG site has a coverage higher or equal to 10x.

**Figure 3:**
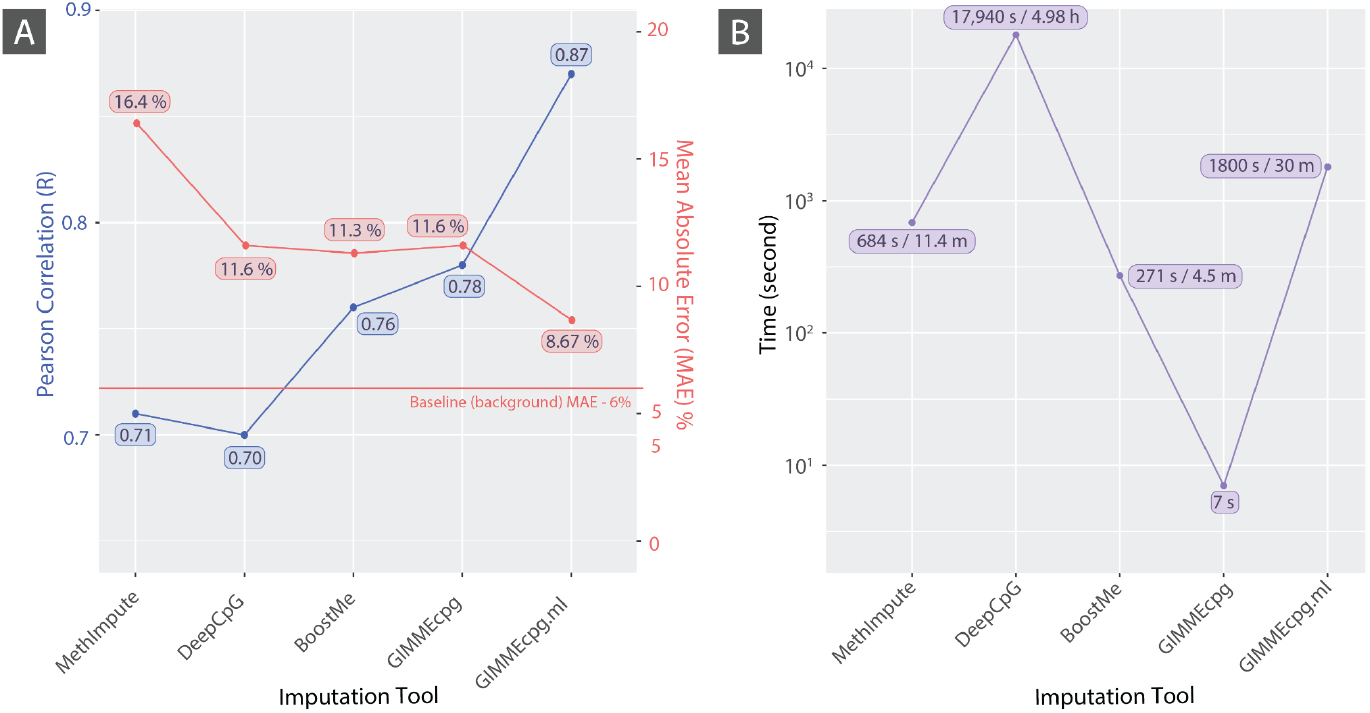
GIMMEcpg.ml Imputation Accuracy at 10x coverage: Accuracy metrics for GIMMEcpg and existing imputation methodologies at 10x coverage level. R: Pearson Correlation; MAE: Mean Absolute Error (%), Time (seconds).

A simplistic imputation approach would be to use the mean to replace missing values. This is particularly appropriate for DNA methylation data because the vast majority of the methylome is highly methylated (and thus close to the mean methylation). In a 10x dataset, it was possible to impute a total of 17.68 million CpG sites with an MAE of 21.9% and an RMSE of 28.2%. As a simple algorithm, the computational resources of such an approach are minimal, with a 2-second runtime using 7GB RAM.

METHimpute, a purpose-built imputation tool designed for DNA methylation data, was more accurate than mean imputation, but at the expense of increased computation complexity. METHimpute took 11 minutes (684 seconds) to impute a total of 17.68 million CpG sites in this 10x downsampled dataset with a Pearson correlation of 0.71 and a MAE of 16.4% and a RMSE of 22.5%.

DeepCpG and BoostMe are purpose-built imputation tools with the best-performing imputation accuracy. For this 10x downsampled dataset, DeepCpg achieved a Pearson Correlation of 0.7 and an MAE of 11.6. However, DeepCpG is extremely slow, taking approximately 5 hours per dataset, despite using a GPU. On the other hand, BoostMe takes roughly 5 minutes to run per dataset with a Pearson Correlation of 0.76 and an MAE of 11.3%.

Interestingly, even though METHimpute had a slightly higher Pearson Correlation than DeepCpG, MethImpute’s MAE was significantly higher (i.e. a difference of 4.8%) than that produced from DeepCpG.

In comparison, GIMMEcpg is roughly 38 times and 2560 times faster than BoostMe and DeepCpG, respectively. Despite being many times faster, GIMMEcpg is roughly just as accurate as BoostMe and DeepCpG. GIMMEcpg has a higher Pearson correlation than BoostMe or DeepCpG while having a similar MAE (i.e. GIMMEcpg at 11.6% vs BoostME at 11.3% and DeepCpG at 11.6%).

In contrast, GIMMEcpg.ml is significantly more accurate than DeepCpG and BoostMe. GIMMEcpg.ml produced a significantly higher Pearson correlation of 0.87 (compared to DeepCpG’s 0.7 and BoostMe’s 0.76) and a significantly lower MAE of 8.67% (compared to DeepCpG’s 11.6% and BoostMe’s 11.3%). This MAE is only 2.67% higher than the baseline MAE, suggesting that GIMMEcpg.ml is able to impute values that are only off by 2.67% compared to the background noise.

As a model learning approach, GIMMEcpg.ml is slower than BoostMe but much faster than DeepCpG. During our internal tests, we were able to reproduce similar accuracies by restricting the analysis time (i.e. to 5 minutes), however, in order to ensure GIMMEcpg.ml was robust, the analysis time was increased to 30 minutes with an early stopping approach used, such that the model stops training once the MAE cannot be further decreased during cross validation.

These benchmarking results demonstrate that GIMMEcpg and GIMMEcpg.ml offer a compelling balance of accuracy and computational efficiency, making them particularly suitable for large-scale methylome studies. To our knowledge, GIMMEcpg is the fastest DNA methylation imputation tool currently available, while GIMMEcpg.ml is the best-performing DNA methylation imputation tool currently available.

### Application to PGP-UK & EpiATLAS datasets

To demonstrate GIMMEcpg’s capability, we applied it to the WGBS datasets from the Personal Genome Project UK (PGP-UK) (Chervova et al. 2019) and the EpiATLAS (EpiATLAS Consortium 2025).

The PGP-UK WGBS data consists of ten WGBS datasets with a coverage of 15.9x. These datasets included 15.9 million CpG sites with a read depth higher than 10x. Running GIMMEcpg took roughly a minute to process all 10 WGBS datasets and imputed an additional 12.2 million CpG sites.

As these WGBS datasets are matched with an Illumina 450k array panel from the same individual, it is possible to assess GIMMEcpg’s accuracy using CpG sites that are common between the array panel and the WGBS dataset. However, it is important to note that differences would be expected due to significantly different processing and analysis technologies between WGBS and array panels.

When considering the entire dataset, the average increase in MAE and RMSE after imputation was 1.4% and 2.7%, respectively. The Pearson correlation decreased by 0.03 on average. When focusing on just the imputed CpG sites, the average increase in MAE and RMSE (compared to baseline error across CpG sites known in both datasets) was 4.1% and 7.2%, respectively.

To demonstrate GIMMEcpg scalability, we applied GIMMEcpg to the EpiATLAS WGBS datasets produced by IHEC (EpiATLAS Consortium 2025). This dataset comprises 645 WGBS datasets of varying coverage. GIMMEcpg took 4 hours and 15 minutes to run on 645 WGBS datasets on a single HPC node (80 CPU threads and 512GB RAM) by sequentially processing each dataset after the next. This included the time to read and process the file and write the output to disk.

Before imputation, the 645 methylomes of the EpiATLAS DNA methylation matrix (minimum 10x read depth for each CpG site) had approximately 6.8 billion missing data points out of a total 18.7 billion data points (29 million haploid CpG sites x 645 datasets). Every single CpG site had a missing data point from at least one sample. Row-wise deletion, where CpG sites with missing data in more than 25% of samples are removed, would cause the number of CpG sites to decrease from roughly 29 million to 9 million. GIMMEcpg was able to impute a total of 2.4 billion data points (using the default 1kb filter), resulting in an additional 20% of data points to the EpiATLAS methylation dataset.

Even though the accuracy of the imputation cannot be determined for these imputed sites within the EpiATLAS dataset, the benchmark within this study helps provide an idea of accuracy based on the dataset’s individual coverage and the corresponding distance to the neighbouring CpG sites.

These applications to real-world datasets demonstrate GIMMEcpg’s efficiency in processing large-scale methylome data and its ability to significantly reduce missing data points, potentially enabling more comprehensive analyses of methylation patterns across samples.

## Discussion

Recent improvements to experimental, instrumental, and computational approaches have made it possible to investigate whole-genome methylomes on an unprecedented scale. Imputation algorithms have greatly helped to improve and harmonise such large-scale datasets but lacked the scalability for routine analysis. To address this limitation and enable all researchers to carry out such large-scale methylome analyses, we have developed GIMMEcpg. In situations where higher accuracy is required, we developed GIMMEcpg.ml.

The evaluation of GIMMEcpg against existing methodologies such as BoostMe and DeepCpG reveals that GIMMEcpg stands out for its exceptional speed while maintaining accuracy that is on par with the more established methods. This speed becomes particularly advantageous when handling the whole methylome datasets that are becoming increasingly common and suggests that efficiency does not necessarily have to come at the cost of precision. Moreover, during our benchmark, existing methodologies (BoostME and DeepCpG) failed to work as effectively at lower coverage datasets (i.e. 5-7x), which is arguably where they would provide the most benefit. In comparison, GIMMEcpg was able to work as effectively at lower coverages.

GIMMEcpg demonstrates that more sophisticated and complex methodologies built using AI or larger training datasets do not always translate to increased accuracy. Interestingly, DeepCpG learns from significantly more information than BoostMe and GIMMEcpg but did not show a proportional increase in accuracy over simpler methods. Similarly, BoostME’s use of a complex gradient boosting algorithm did not improve accuracy compared to GIMMEcpg. In fact, the increased input dataset size and complex methodologies resulted in drastically increasing the computational requirements as well as the runtime. Therefore, this highlights the importance of benchmarking against more traditional and less complex methodologies.

Furthermore, GIMMEcpg.ml significantly surpasses the accuracy of both BoostMe and DeepCpG and is expected to be ideal for cases where higher accuracy is required at the expense of slower runtime - for example, when identifying DMPs with small differential methylation, the imputation accuracy becomes more critical. GIMMEcpg.ml is therefore a favourable alternative to DeepCpG and BoostMe for most applications. However, when additional data types (such as histone modifications or RNA-seq) are available, tools like ChromImpute should be explored. Nonetheless, most routine DNA methylation studies do not include multiple epigenomic modalities and thus cannot be used with ChromImpute. Additionally, there isn’t always enough biosample available for generating this multi-dimensional data.

The increasing production of DNA methylation datasets, fueled by advances in sequencing technologies and national biobanks, points to a future where the efficiency and scalability of computational tools will be even more critical. National Biobanks, such as in Estonia and Japan have started to routinely produce whole methylome datasets at scale and the UK Biobank recently announced to generate 50,000 methylomes using Nanopore-seq (Anon n.d.). Furthermore, technologies like Nanopore-, SMRT, TAB- and Biomodal-seq are able to produce methylome datasets at no or little additional cost, making them even more accessible to the scientific community. Given these advances, it can be expected that the number of methylome datasets will grow exponentially over the coming years, requiring algorithms like GIMMEcpg to enable swift analysis at scale without compromising accuracy.

It’s worth noting that GIMMEcpg is expected to become even faster as the underlying technology (https://pola.rs/) is under active development with a large community. The technology has recently added support for GPUs and can automatically be used (if available) for relevant parts of the underlying algorithm where appropriate, further improving its performance.

GIMMEcpg and GIMMEcpg.ml are available as docker images and can thus be quickly deployed to cloud computing environments. Furthermore, the drastically reduced computational requirements of GIMMEcpg compared to alternative tools and faster run times would also equate to drastically lower cloud computing costs. The reduced computational load would also likely equate to a decreased carbon footprint (Grealey et al. 2022).

Despite its advantages, GIMMEcpg does have some limitations. Due to its reliance on neighbouring information to impute missing sites, its application is mainly limited to data where neighbouring CpG sites are close enough to the missing site to have highly correlated methylation values. In general, this means that the neighbouring CpG and missing sites should be within 1kb of each other.

Since the feature dataset for GIMMEcpg is built using only the information within each sample, GIMMEcpg cannot integrate information from other data types, such as RNA-Seq or histone data. Similarly, the imputation model for GIMMEcpg is built on a per-sample basis and does not translate across samples within a cohort or to different cell types. Furthermore, GIMMEcpg was developed for bulk WGBS data, and its utility on single-cell methylation data has not been confirmed.

In conclusion, GIMMEcpg and GIMMEcpg.ml represents a significant advancement in methylome imputation, offering a balance of speed, accuracy, and scalability that is well-suited to large-scale methylome studies. GIMMEcpg and GIMMEcpg.ml are easily accessible through Python and R libraries, and its performance and efficiency make it a valuable tool for researchers working with whole-genome methylation data.

## Methods

### Datasets

The datasets used for the benchmark carried out in this work have been produced by IHEC and are part of the EpiATLAS (EpiATLAS Consortium 2025). Two deeply sequenced WGBS (IHECRE00000101.3 and IHECRE00000155.3) were produced from CD14-positive, CD16-negative classical monocytes. IHECRE00000101.3 was produced from venous blood from a healthy, 60-65 years old female donor (C004SQ), while IHECRE00000155.3 was produced from cord blood from a healthy, 0-5 years old female donor (C005PS).

These WGBS datasets were sequenced using purified cells on an Illumina HiSeq 2000 instrument. The datasets were prepared using 101 base pair, pair-ended reads. To achieve deep coverage, a total of 11 (IHECRE00000101.3) and 10 (IHECRE00000155.3) sequencing runs were carried out for each sample, and the sequenced reads were then merged before further processing.

The WGBS summary CpG count files for IHECRE00000101.3 and IHECRE00000155.3 are available through IHEC, while the raw data is available on EGA (EGAX00001097774 and EGAX00001097775 respectively).

### Downsampling

These two deeply sequenced datasets have a CpG coverage of approximately 100x. This is more than three times the standard coverage (30x) that is recommended by IHEC for WGBS. As such, this makes these two datasets ideal for producing multiple downsampled datasets, each with a lower CpG coverage.

To downsample these two datasets, a random read (together with its paired mate) is selected from all mapped reads (i.e., from the BAM file) and then merged to produce another dataset. A new mapped BAM file with a lower sequencing coverage can be produced *in silico* by selecting a specific percentage of read pairs.

The samtools view command from the Samtools package (version: 1.11) was used with the ‘-s’ parameter to randomly select random read pairs from the input BAM file, producing a smaller, downsampled BAM file. Moreover, to increase the randomness of which read pairs are selected, a different seed number was used each time the input file was downsampled. The downsampled BAM file can then be processed using the standard pipeline to extract the downsampled dataset’s methylation information.

A total of 24 datasets were downsampled for each dataset, with three datasets produced at each of the following eight different coverage levels: 5%, 7%, 10%, 15%, 20%, 25%, 30% and 60%. To ensure a high level of diversity between the 24 datasets, a different seed number was used each time data was downsampled. In conclusion, a total of 48 downsampled datasets were produced for both samples (i.e. 24 datasets for each sample) by using three different seed numbers. A summary of the downsampled datasets is shown in Supplementary Table 1.

### GIMMEcpg R and Python Packages

GIMMEcpg is available as both R and Python packages. Both implementations use the Polars library and its lazy application programming interface (API) (Vink et al. 2024). These features allow GIMMEcpg to process data in parallel and carry out automatic query optimisation. In addition, GIMMEcpg has been built with an option to handle large out-of-RAM datasets by utilising the lazy API’s streaming mode. Before imputing, GIMMEcpg first collapses CpG sites on complementary strands into one row. Following this, GIMMEcpg filters out CpG sites with low read coverage; any sites not meeting the minimum coverage threshold will be considered missing and subsequently imputed. The default coverage threshold is ten reads. GIMMEcpg then cross-references the sample methylation data with a reference containing all possible CG sites in the human genome to identify missing CpG sites for imputation. The package includes two reference files, one each for hg38 and hg19. Users may also choose to supply their own reference files or generate them with the provided R script.

Only autosomal CpG sites are considered for constructing the feature dataset and imputing missing methylation values. In addition, any CpG sites falling within the ENCODE blacklisted regions are excluded from both the feature dataset and imputation (Amemiya et al., 2019). Exclusion of ENCODE blacklisted CpG sites is the default behaviour; however, it is possible to override it or to provide a custom blacklist.

GIMMEcpg depends almost entirely on Polars to perform the required calculations. On the other hand, GIMMEcpg.ml relies on H2OAutoML’s supervised machine learning algorithm to train several models and stacked ensembles based on the known CpG sites in the dataset (LeDell & Poirier 2020). The models are then ranked, and the best model based on mean residual deviance is used to impute the missing methylation values. If a maximum neighbour distance has been specified, GIMMEcpg will only use known CpG sites within the neighbouring distance threshold for the feature set and impute only the missing sites that are less than the specified distance from its immediate neighbours. The default threshold for maximum neighbour distance in both GIMMEcpg and GIMMEcpg.ml modes is 1kb.

## Notes

### Competing Interest Statement

The authors have declared no competing interest.

