## Supplementary Figures for "GIMMEcpg: Global Imputation of Mean CpG MEthylation in Real-time"

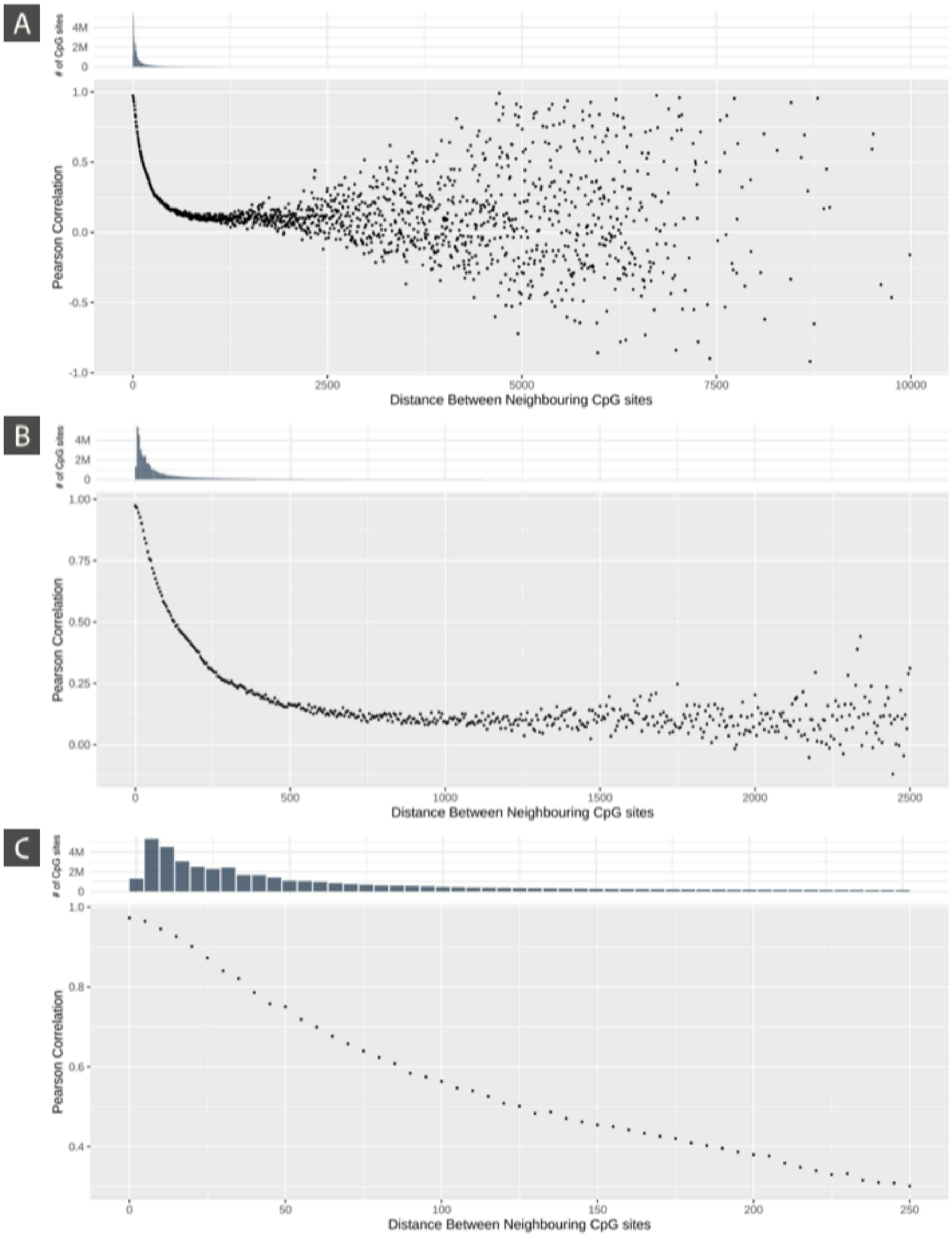


***Supplementary Figure 1: Pearson correlation between neighbouring CpG sites.*** *The methylation value and distance between a CpG site and its immediately upstream neighbouring CpG site were extracted from two high-quality (100x) bulk-cell WGBS data from the International Human Epigenome Consortium (IHEC). Next, neighbouring CpG sites were binned based on a 5bp neighbouring window and the Pearson correlation was calculated using all the CpG site Pairs (i.e. neighbouring CpGs) falling*

*within the window (i.e. all neighbouring CpG sites that are between 30-35bp from one another are binned together and the Pearson Correlation calculated between the CpG pair). Panels B and C are a zoomed-in version (along the x-axis) of panel A to show more detail.*

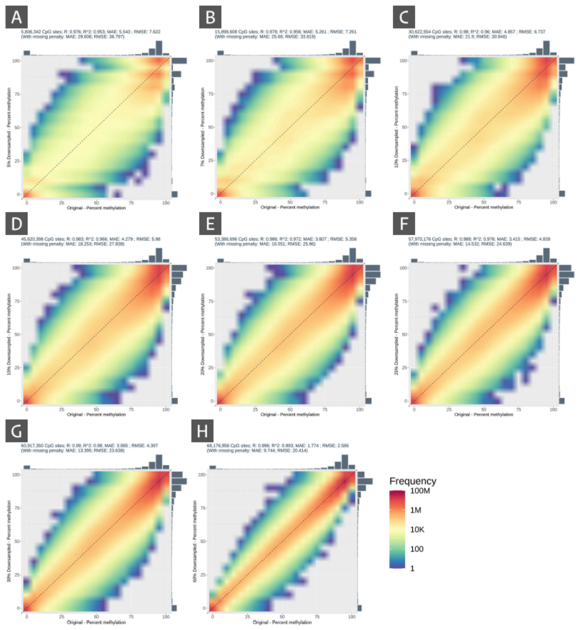


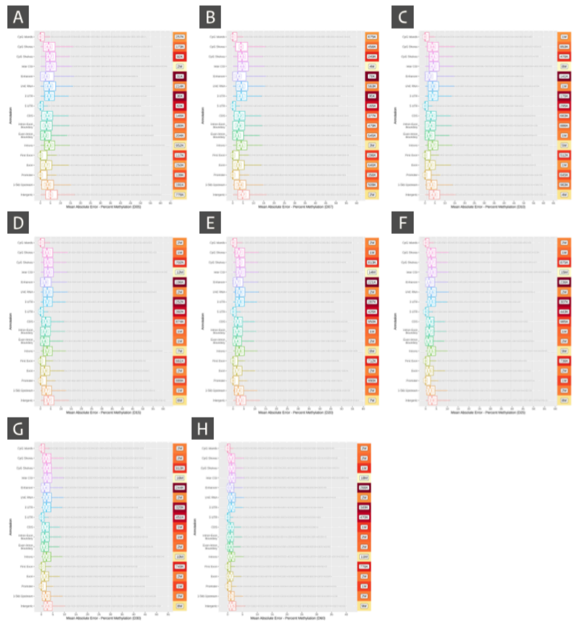


***Supplementary Figure 2: Baseline error in the downsampled datasets*** *(i.e. residual noise in the dataset due to being at lower coverage). Distributions comparing the methylation value in the downsampled dataset and the original dataset's corresponding actual methylation value. Each panel only includes information from CpG sites with a greater than 10x read depth. Any deviation from the y = x-axis corresponds to an error in the methylation value. Each panel corresponds to a specific coverage level (i.e. panels A-H correspond to 5x, 7x, 10x, 15x, 20x, 25x, 30x and 60x, respectively); also shown in the y-axis title. Each panel includes summarised information from six downsampled datasets at that specific coverage level.*

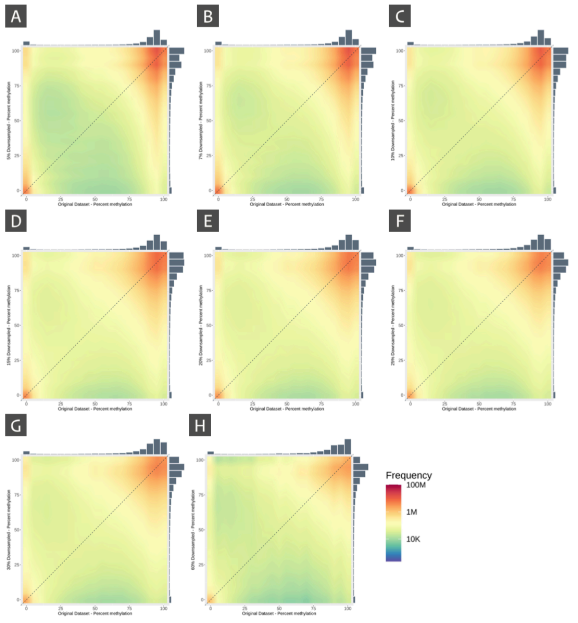


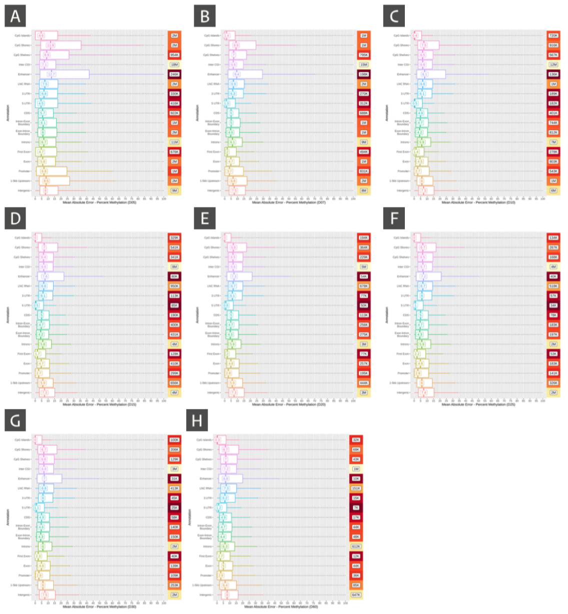

***Supplementary Figure 3: A] GIMMEcpg Imputation:*** *Methylation distribution of imputed values, comparing the imputed methylation value and the corresponding actual methylation value in the original dataset. Each panel only includes information from CpG sites with a greater than 10x coverage. Any deviation from the y=x axis corresponds to an error in the methylation value. Each panel corresponds to a specific coverage level (i.e. panels A-H correspond to 5x, 7x, 10x, 15x, 20x, 25x, 30x and 60x, respectively), shown in the y-axis title. Each panel includes summarised information from six downsampled datasets at that specific coverage level. However, it is important to note that these plots do not take into account the baseline error and consequently, are expected to be overestimated.* ***B: GIMMEcpg Imputation:*** *Box plots showing the distribution of the absolute error in the imputed value, separated based on the genomic annotation of the CpG site. Outliers are not shown on the plot due to the number of outliers; rather, the box-plot whiskers are extended to the maxima and minima with a dashed line. Each panel corresponds to a specific coverage level (i.e. panels A-H correspond to D05, D07, D10, D15, D20, D25, D30 and D60, respectively), shown in the y-axis title. Each panel includes summarised information from six downsampled datasets at that specific coverage level. The number of CpG sites in each genomic annotation type is highlighted in the colour scale on the right of each panel (with the red colour denoting a lower number of CpG sites).*

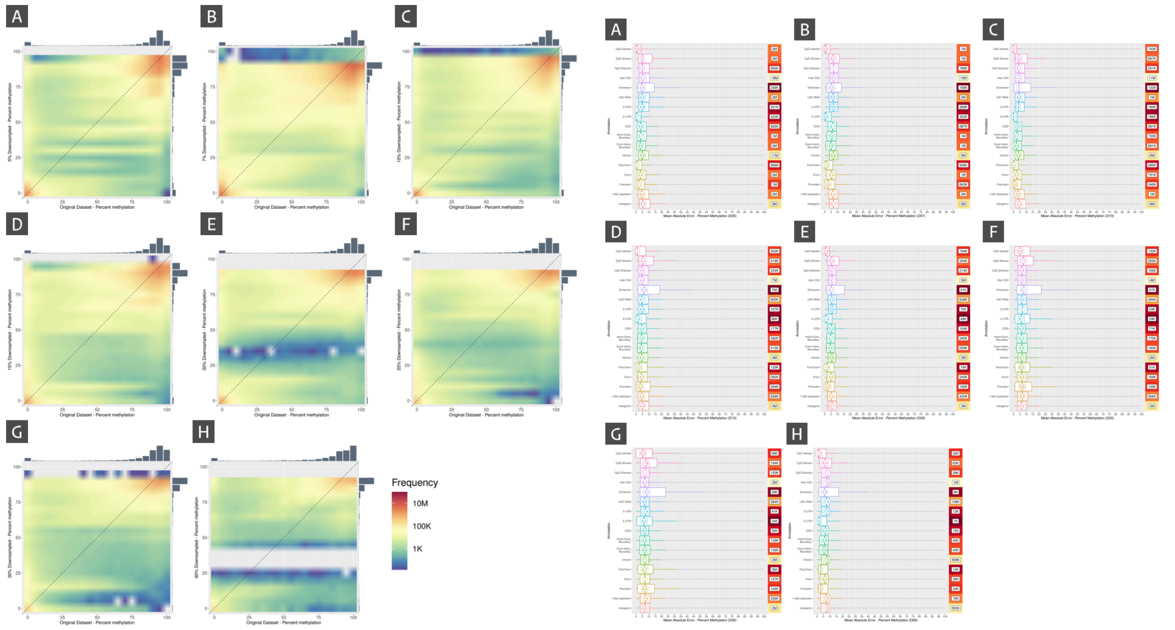


***Supplementary Figure 4: [A] GIMMEcpg.ml Imputation****: Methylation distribution of imputed values, comparing the imputed methylation value and the corresponding actual methylation value in the original dataset. Each panel only includes information from CpG sites with a greater than 10x coverage. Any deviation from the y = x-axis corresponds to an error in the methylation value. Each panel corresponds to a specific coverage level (i.e. panels A-H correspond to D05, D07, D10, D15, D20, D25, D30 and D60, respectively), shown in the y-axis title. Each panel includes summarised information from six downsampled datasets at that specific coverage level.* ***[B] GIMMEcpg.ml Imputation:*** *Box plots showing the distribution of the absolute error in the imputed value, separated based on the genomic annotation of the CpG site. Outliers are not shown on the plot due to the number of outliers; rather, the box-plot whiskers are extended to the maxima and minima with a dashed line. Each panel corresponds to a specific coverage level (i.e. panels A-H correspond to D05, D07, D10, D15, D20, D25, D30 and D60, respectively), shown in the y-axis title. Each panel includes summarised information from six downsampled datasets at that specific coverage level. The number of CpG sites in each genomic annotation type is highlighted in the colour scale on the right of each panel (with the red colour denoting a lower number of CpG sites).*
